# Transcranial Photobiomodulation on the Left Prefrontal Cortex Enhances Mandarin Chinese L1 and L2 Complex Sentence Processing Performances

**DOI:** 10.1101/2024.01.22.576680

**Authors:** Mingchuan Yang, Yang Liu, Zhaoqian Yue, Guang Yang, Xu Jiang, Yimin Cai, Yuqi Zhang, Xiujie Yang, Dongwei Li, Luyao Chen

## Abstract

This study investigated the causal effect of transcranial photobiomodulation (tPBM) over the left prefrontal cortex (LPFC) on syntactically complex Mandarin Chinese first language (L1) and second language (L2) sentence processing performances. Two (L1 and L2) groups of participants (thirty per group) were recruited to receive the double-blind, sham-controlled tPBM intervention, followed by the sentence processing, the verbal working memory (WM), and the visual WM tasks. Results revealed a consistent pattern for both groups: (a) tPBM enhanced sentence processing performance but not verbal WM and visual WM performance; (b) Participants with lower sentence processing performances under sham tPBM benefited more from active tPBM. Taken together, the current study substantiated that tPBM enhanced L1 and L2 sentence processing ability directly without verbal WM interference, and would serve as a promising and cost-effective noninvasive brain stimulation (NIBS) tool for future applications on upregulating the human language faculty.

**Highlights:**

a. The first study that applies tPBM to (complex) sentence processing.
b. tPBM enhances sentence processing performances in both Mandarin L1 & L2 speakers.
c. tPBM directly enhances sentence processing without the interference of verbal WM.
d. A causal role of LPFC for sentence processing through active tPBM.
e. Opening up the promising application prospect for tPBM on sentence processing.

## 1 Introduction

The competence in processing sentences especially with complex syntactic structures is a hallmark of human high-level cognition and is viewed as the core of human language faculty (Dehaene et al., 2015; Fitch, 2014; Friederici, 2017; Goucha et al., 2017; Hauser et al., 2002; Nelson et al., 2017). With the development of neurolinguistics, how the brain processes language has been extensively investigated. Several brain regions have been proved to engage in sentence processing [such as the left prefrontal cortex (LPFC; Friederici et al., 2006b; Makuuchi et al., 2009; Meyer et al., 2012; Santi & Grodzinsky, 2010; Xu et al., 2020) and the left posterior temporal cortex (LpTC; Ben-Shachar et al., 2004; Friederici et al., 2009; Goucha & Friederici, 2015; Kinno et al., 2008; Obleser & Kotz, 2010)] and to manifest their functional or anatomical plasticity across various kinds of participants [such as healthy participants & patients (Barbier et al., 2019; Ilves et al., 2014; Thompson, 2019; Thompson et al., 2021), young adults & elder adults (Mueller, 2009; Wingfield & Grossman, 2006), adults & children (Davidson, 2010; Müller et al., 1999), first language (L1) speakers & second language (L2) speakers (Davidson & Indefrey, 2009; Proverbio et al., 2002; Steinhauer & Kasparian, 2020; Wang P. et al., 2021, 2022, Wei et al., 2024)].

Considering the significance of sentence processing and for the sake of improving its abilities as well as relieving dysfunctions, intervention towards sentence processing ability has already been followed with great interest for decades. Owing to the feasibility of shaping the brain, the noninvasive brain stimulation (NIBS), which causes electrophysiological or metabolic effects through physical or chemical approaches to alter brain activities, has become a promising method of modulation towards language ability (Hussey et al., 2015; Minamoto et al., 2014; Ohn et al., 2008; van der Burght et al., 2023). Not only towards patients with language ability deficiency to restore the affected functions (Cotelli et al., 2011; Hartwigsen & Siebner, 2013; Thiel et al., 2013), NIBS is also expected to be applied on healthy adults with continual neural plasticity. Although adults’ language network is fully mature in both structure and function, it also appears to remain plastic during the whole lifespan in the course that we continue to learn and process various kinds of language information (either in L1 or L2; Li et al., 2014; Schlegel et al., 2012; Stein et al., 2012; Wang P. et al., 2021). Moreover, healthy adults with relatively small individual variance compared to patients can serve as an ideal case to explore NIBS’s modulatory effects (Hartwigsen et al., 2013; Qu Xin. et al., 2022). Therefore, the investigation of NIBS’s effects on L1 and L2 speakers’ sentence processing ability holds great significance and is supposed to arouse particular attention.

### 1.1 The neuromodulation through tPBM

Drawn on the technique of NIBS, the effect of neuromodulation on language ability has been explored in the past decades mainly using transcranial electrical stimulation (tES) and transcranial magnetic stimulation (TMS) (Cattaneo et al., 2011; Fertonani et al., 2010; Holland et al., 2011). It is worth noting that a large body of tES and TMS studies were interested in the explorations of the causal relationships between the target regions and the behavioral/neural changes in healthy participants by utilizing inhibitory protocols (e.g., Sakreida et al., 2019; Ware et al., 2021; Zhu & Snowman, 2020) while leaving the facilitatory/enhancement effects underspecified. A recent meta-analysis also pointed out that the modulation effectiveness of TMS on specific aspects of language ability (e.g., syntactic ability) was relatively limited (Qu Xin. et al., 2022). Therefore, it is necessary to apply an alternative technique with a higher availability of enhancement effect—transcranial photobiomodulation (tPBM)—to upregulate language ability. The tPBM is a newly-developed NIBS technique and can regulate mitochondrial respiration and cellular functions by shining red-to-near-infrared light (600– 1100 nm) on the cerebral cortex through the cranium, in a nondestructive and nonthermal optical fashion, and specifically, the photochemical reaction within brain cells rests on that complex Ⅳ of the mitochondrial respiratory chain is upregulated by absorbing photonic energy to modulate cytochrome c oxidase (CCO), which results in the increased oxygen consumption and adenosine triphosphate (ATP) formation (Barrett & Gonzalez-lima, 2013; Eells et al., 2004; Tian et al., 2016; Urquhart et al., 2020; Wang X. et al., 2022; Wong-Riley et al., 2005). Since brain physiology is dependent on oxygenation for energy utilization, tPBM can finally boost brain cognition (Lee et al., 2023; Wang X. et al., 2022; Zhao et al., 2022; see Wang X. et al., 2022 for more detailed information about tPBM functional mechanism).

Recently, tPBM applied on the human forehead has been evidenced to modulate the prefrontal cortex (PFC) by improving the PFC-based cognitive functions in healthy adults, elderly people, or patients with psychiatric and neurological disorders (Chao et al., 2020; Kerppers et al., 2020; Qu Xiu. et al., 2022; Zhao et al., 2022; see Lee et al., 2023 for a systematic review). The beneficial effect was found most robustly on PFC-modulated memory ability (Barrett & Gonzalez-lima, 2013; Chan et al., 2021; Holmes et al., 2019; Hwang et al., 2016; Qu Xiu. et al., 2022; Zhao et al., 2022). Barrett and Gonzalez-lima (2013) first conducted a controlled study demonstrating that the performance on a delayed match-to-sample (DMS) memory task was improved for tPBM stimulated group as opposed to the control (placebo) group. Zhao et al. (2022) found that 1064-nm tPBM on the right PFC significantly enhanced the visual working memory (WM) capacities in healthy adults and proposed the mediating effect of electrophysiological activities. In addition, tPBM also produced enhancement for attention, executive functions, and other PFC-based abilities according to recent studies (Blanco et al., 2017; Chan et al., 2019; Moghadam et al., 2017; see also Lee et al., 2023 and Salehpour et al., 2019 for systematic review and meta-analysis).

### 1.2 PFC engagement in sentence processing

When it comes to the cognitive functions of the prefrontal cortex, it is inevitable to involve the critical high-level sentence processing competence (e.g., Friederici et al., 2003, 2006b, 2017; Jeon, 2014; Hagoort, 2013; Malik-Moraleda et al., 2022; Martins et al., 2019; Vigneau et al., 2006). The processing of sentences is proposed to be based on the fundamental operation of *merge*, a process defined by the Generative Linguistics to combine two syntactic objects into a larger new constituent (Chomsky, 1995; Friederici, 2017; Miyagawa et al., 2013; Zaccarella & Friederici, 2015). Such a computational ability to build up the syntactic hierarchies is believed to play an essential role in human language faculty, which was found to be largely dependent on the functions of the left inferior frontal gyrus (LIFG) within the ventral part of LPFC (e.g., Chen et al., 2021a, 2023; Friederici, 2017; Friederici et al., 2006b; Jeon, 2014; Liu et al., 2023; Makuuchi et al., 2009; Meyer et al., 2012; Santi & Grodzinsky, 2010; Zaccarella et al., 2015, 2017a; see also Zaccarella et al., 2017b for the meta-analysis on the neurobiology of merge). Zaccarella et al. (2015) provided evidence that Brodmann Area (BA) 44, a relatively posterior part of LIFG, played the primary supporting role for merge when processing syntactic phrases compared to word-list sequences. In addition, the activation of LIFG directly correlates with the syntactic complexity as shown by the studies focusing on the processing of noncanonical sentences involving word scrambling, syntactic movement, and multiple syntactic embeddings (e.g., Ben-Shachar et al., 2004; Caplan et al., 2008; Friederici et al., 2006b; Makuuchi et al., 2009, 2013; Meyer et al., 2012; Röder et al., 2002; Santi & Grodzinsky, 2010). Evidence of artificial hierarchical grammar processing from Chen et al. (2021a) and Liu et al. (2023) further indicated that LIFG engages in the build-up process of syntactic hierarchies. In particular, for the first time, Liu et al. (2023) found a significant correlation between the signal intensity of the relatively posterior part of LIFG as identified in their artificial merge grammar processing and the behavioral performances of natural complex sentence processing. These studies, thus, converge on and underly the critical role of LIFG within the ventral LPFC in merge during sentence processing as a syntactic engine.

### 1.3 Hypothesis of two pathways of tPBM effects on sentence processing

From the perspective of LPFC’s (esp., LIFG’s) functions in sentence processing, it is of great significance to test whether tPBM on LPFC could exert enhancement on sentence processing performances in the current context of few explorations of tPBM on human language faculty. Nevertheless, the processing of sentences (esp., complex sentences) would inevitably maintain a number of sentence components active in verbal WM until the construction of syntactic structure as well as the integration of syntactic and semantic information are completed (Santi & Grodzinsky, 2007). The highly interactive relationship between sentence processing and verbal WM has been certified by a myriad of studies (e.g., King & Kutas, 1995; King & Just, 1991; Makuuchi et al, 2009; Vos et al., 2001). The correlation between verbal WM span and syntactic computational ability was found in both behavioral and neurophysiological evidence (Just & Carpenter, 1992; McDonald, 2006; Vos et al., 2001; Fiebach et al., 2004). Moreover, results from multiple neural localization studies emphasize the engagement of LPFC underlying verbal WM functions (Fregni et al., 2005; Ohn et al., 2008; Smith & Jonides, 1998). A further study (Makuuchi et al., 2009) proposed a picture in which syntactic merge represented in the left pars opercularis (LPO) was segregated but highly interconnected with the syntax-specific memory-related profile housed in the left inferior frontal sulcus (LIFS). When it comes to the evidence that tPBM did benefit the WM ability, two possible functional pathways of tPBM effect on sentence processing might emerge: one hypothesizes that tPBM modulated sentence processing directly, independent of the WM capacity, and the other assumes that the effect of tPBM on sentence processing should be interfered by the verbal WM to a certain extent (Figure 1). Therefore, whether the tPBM on the LPFC (esp., LIFG) would upregulate the sentence processing performances independently awaited to be explored.

**Figure 1.**
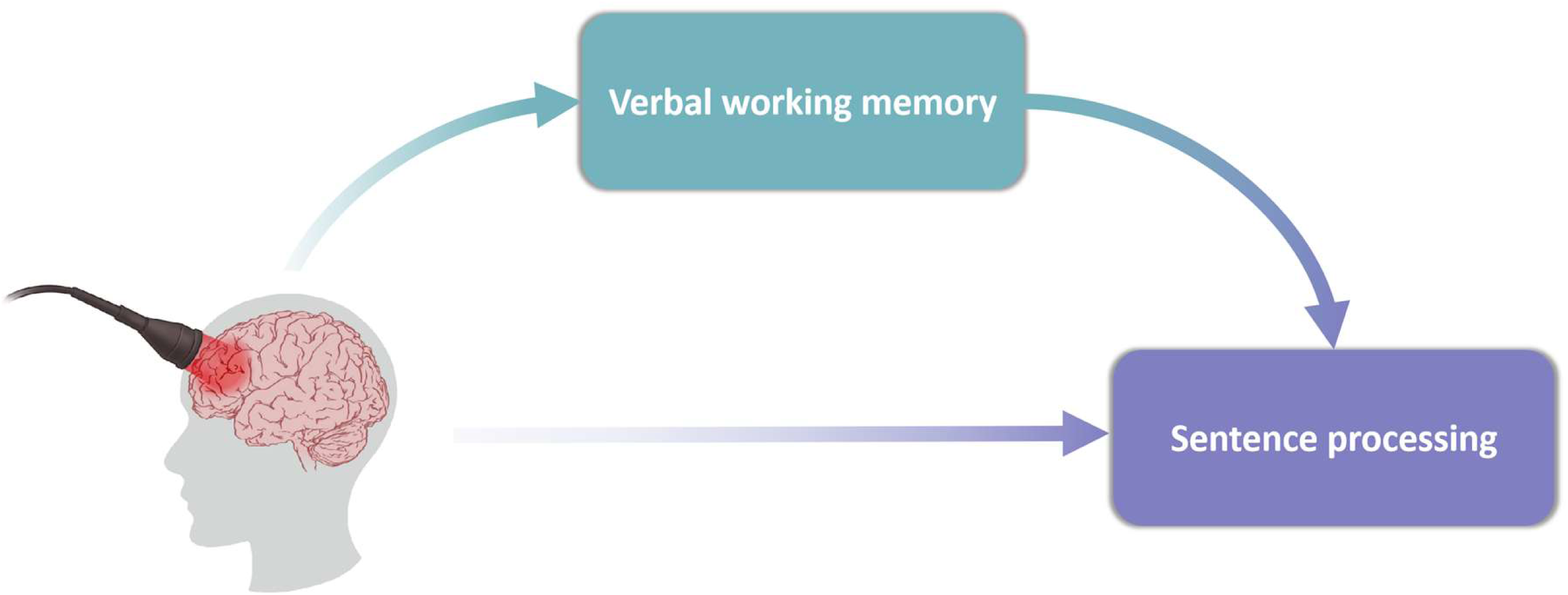
Two possible functional pathways of tPBM effect on sentence processing: the lower one hypothesizes that tPBM modulated sentence processing directly, independent of the working memory (WM) capacity; the upper one assumes that the effect of tPBM on sentence processing should be interfered by the verbal WM.

### 1.4 A developmental view of tPBM effects on sentence processing

In order to detect the tPBM effect on sentence processing, healthy adults who are native speakers of the target language with relatively small individual variance (compared to L2 learners with higher internal variance considering their differed language background, L2 proficiency level, age of acquisition, etc.) can serve as an ideal case and a starting point for the initial exploration (Hartwigsen et al., 2013; Qu Xin. et al., 2022). Also, evidence showing the large plasticity of L1 (Wang P. et al., 2021, 2022) speakers underlie the feasibility of intervention towards the language ability on them.

Moreover, investigating tPBM effects on sentence processing ability from a language developmental view is also of our primary interest, which could further guide applications of tPBM on groups struggling with language ability deficiency in the near future. Among a wide range of people facing problems with sentence processing, L2 learners who have normal non-language ability (e.g., attention and executive function; compared to patients) enable us to make further investigations, by which the confounding effects brought by the non-language factors could be controlled to a relatively low extent. From recent studies, L2 learners were found to process sentences also with LPFC (esp., LIFG) highly involved, which suggested that L1 and L2 speakers share a common brain area in LPFC to accomplish sentence processing (e.g., Chen et al., 2019, 2021b; Golestani et al., 2006; Jeon & Frederici, 2013; Mueller et al., 2014; Nakagawa et al., 2022; Nachi & Sakai, 2009; Sakai et al., 2009; Tao et al., 2021; Umejima et al., 2021; Wartenburger et al., 2003). Specifically, Chen et al. (2019, 2021b) proposed that native Korean speakers showed significant activation in posterior LIFG when reading artificial sentences generated by the Chinese-like grammar based on word category information. Wartenburger et al. (2003) found that the late bilinguals induced greater activation when processing L2 sentences in LPFC than the early ones and even than when they processed L1. A recent study on Japanese English learners (Nakagawa et al., 2022) dissociated the brain areas responsible for semantic from syntactic processing and pointed out that LIFG is involved in grammatical encoding in the process of phrase production. Meanwhile, a study of NIBS revealed that L2 learners’ ability of syntactic processing showed plasticity and could be enhanced through stimulating LIFG (de Vries et al., 2010). Therefore, it is reasonable to hypothesize that tPBM on LPFC could show enhancement effects on sentence processing for L2 learners as well.

Furthermore, when it comes to the two hypothesized effective pathways of tPBM as mentioned in Section 1.3, questions appear pronounced whether L1 speakers and L2 learners would show parallel or divergent patterns of the tPBM effect on complex sentence processing. One may predict that L1 speakers and L2 learners might differ in the effective pathway of tPBM on sentence processing (see Section 1.3). The WM for L2 elements (i.e., the WM to hold various language information of L2 in mind) is less efficient and its ability is obviously worse than the homologue of L1 speakers, such that the sentence processing in L2 demands WM to a larger extent (Ardila, 2003; McDonald, 2006).

### 1.5 The present study

This study aimed to explore the tPBM enhancement effects on complex sentence processing in both L1 and L2 participants after the stimulation on LPFC. Complex sentences with relative clauses (RC) embedded are challenging even for the L1 healthy adults and were, therefore, used as sentence processing materials in the current study (see also Wang P. et al., 2021, 2022). Meanwhile, to test the aforementioned two-pathway hypothesis of tPBM effects, a verbal WM task was also developed in the present study. Additionally, to test whether tPBM effects on LPFC are domain-specific, a visual WM task, which has already been certified unrelated to LPFC (Zhao et al., 2022), was manipulated as a control task in the current study. By recruiting Mandarin Chinese L1 speakers and L2 learners, the present study investigates the following questions:

a. Can tPBM on LPFC facilitate sentence processing?
b. If the answer to question (a) is yes, whether the effect of tPBM directly applies to sentence processing or with the modulation on verbal WM as an intervention?
c. What is the relationship of the tPBM effective pathway (see Section 1.3) between L1 and L2 groups?

Answers to these questions could be instructive to the utilization of tPBM on the upregulation of sentence processing and provide profound insights into the functional neural plasticity of L1 and L2 sentence processes.

### Data availability

The data files, lists of materials, and codes of data analyses can be downloaded via the Open Science Framework at https://osf.io/e35ac/.

## 2 Methods

### 2.1 Participants

Thirty Mandarin Chinese native speakers (14 males, 22.47 ± 1.74 years) and thirty Mandarin Chinese L2 learners whose native languages were Thai or Vietnamese (7 males, 21.97 ± 2.92 years; 6 Thai and 24 Vietnamese) participated in the current study. Thai and Vietnamese are both head-initial languages with postnominal RC locations with regard to the language typology, which mirrors the order of relative clause and head noun in Chinese^1^ (Chu, 2020; Liu, 2019; Mao, 2018). Therefore, we recruited these participants from similar language backgrounds under the perspective of complex sentence processing. All Mandarin L2 speakers were overseas students studying in mainland China during the sessions of experiment, whose Chinese proficiency had reached the intermediate or advanced level of the HSK (i.e., Hanyu Shuiping Kaoshi, a standardized Chinese proficiency test, ranging from bands 1 with low proficiency to 6 with advanced proficiency) band-4 or above. All L2 participants completed a questionnaire of language background additionally. They began to learn Chinese as a second language at an average age of 16.25 ± 4.19 years and the mean length of learning was 5.32 ± 3.67 years. They all reported Mandarin Chinese as the second most familiarized language after their mother tongues.

All participants were right-handed, with normal or corrected-to-normal vision and no color blindness or color weakness. None of them reported reading difficulty or any history of psychiatric or neurological diseases. They all signed the consent prior to the experiment and received a monetary reward afterward. This study was approved by the Ethics Committee of Beijing Normal University. Data from four participants (two L1 participants and two L2 participants) were excluded because of the relatively lower data quality (i.e., more than 20% of the trials were missed for not pressing keys on the keyboard) or of the unaccomplishment of the whole experiment. Therefore, twenty-eight L1 participants’ (13 males, 22.36 ± 1.73 years) and twenty-eight L2 participants’ (6 males, 22.00 ± 2.99 years) data remained as valid and were entered into subsequent formal analyses.

### 2.2 Materials

Materials for sentence processing task, verbal WM task, and visual WM task were prepared respectively. The detailed settings of experimental materials for each task were delineated as follows.

#### Sentence processing materials

Syntactically complex sentences containing relative clause (RC) were adopted for sentence processing task. RC is a kind of subordinate clause that modifies a head noun and is embedded within a noun phrase. In Chinese, a language with head-final RC pattern, noun phrase containing RC has a structure of “inflection phrase + De (的, complementizer) + head noun”. For example, in “支持花花的小孙帮助老张 (literal glosses: support Huahua de Xiaosun know Laozhang; translation: Xiaosun who supports Huahua helps Laozhang)”, “支持花花的小孙” is a noun phrase with a RC of “支持花花的”. “小孙” is extracted from the clause and leaves a gap. “小孙” is coindexed with the gap and is called the filler because it should fill the gap (Figure 2A). To comprehend this kind of sentences needs reordering and integration across a long-distance filler-gap dependency, necessary for hierarchical syntactic building. Thus, sentences with RCs contain great complexity of syntactic computation, the processing of which highly involves LIFG (Santi & Grodzinsky, 2010; Xu et al., 2020) and is assumed by the present study to show high potential to be modulated by tPBM on LPFC.

**Figure 2.**
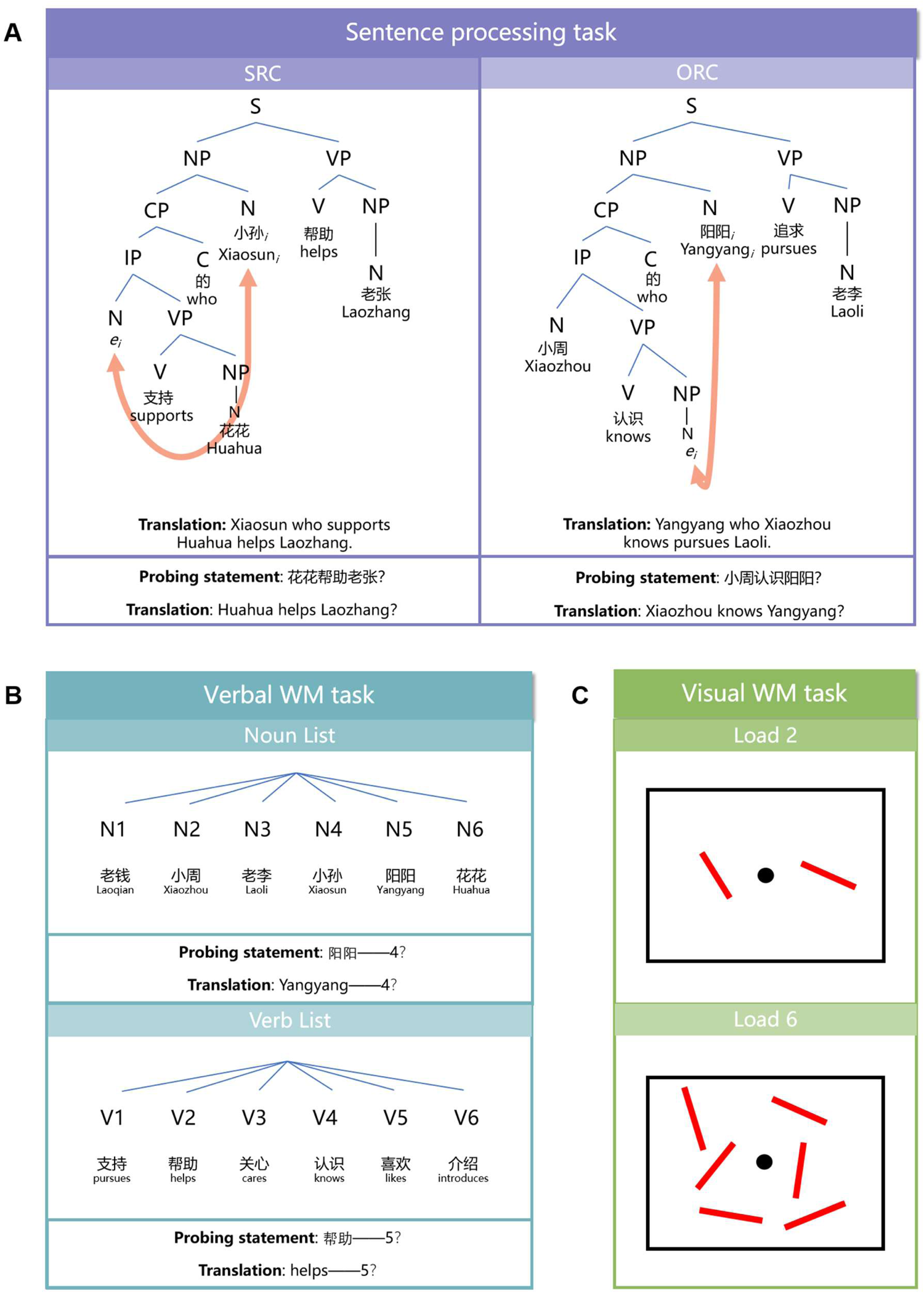
**(A)** Examples of materials in the sentence processing task. Every sentence contains either an SRC or an ORC with subject or object being extracted and leaving a gap. Sentence structures are presented in the form of syntactic tree. Every Chinese word is attached to its English literal gloss and the English translation of the whole sentence and probing statement are provided below. The gap (*e*) and the target dependent noun (N) are co-indexed by the subscript “*i*” and linked by an orange arc. S: subject; NP: noun phrase; N: noun; VP: verb phrase; V: verb; CP: complementizer phrase; IP: inflection phrase; C: complementizer; *e*: gap. **(B)** Examples of noun (name in Chinese) word lists and verb word lists in verbal WM task. Each list consists of 6 words in linear sequence. Each word in word lists is attached with its English literal gloss. The probing statement and its English translation are presented below. **(C)** Examples of materials in visual WM task. A fixation point is surrounded by two and six bars in the condition of load 2 and load 6.

In order to increase the variation of the materials, a total of 72 complex sentences containing RCs were generated, including 36 sentences with subjective relative clauses (SRC) and 36 sentences with objective relative clauses (ORC) embedded at either subject or object positions of the main clauses. Specifically, 12 two-syllable verbs selected from HSK-4 vocabulary syllabus and 12 two-syllable common names (i.e., nouns) from HSK textbooks were used to build all complex sentences. Moreover, word frequencies and the frequencies of collocation between two nouns/verbs or a noun and a verb were carefully controlled so that participants were unable to process the sentences or make judgements with any possible strategies unrelated to language processing. Following Xu et al. (2020) and Liu et al. (2023), a probing statement of thematic relation (i.e., the relation of “who did what to whom”) was attached to each sentence trail for the correctness judgement to detect participants’ performance of syntactic processing (Figure 2A). The probing sentences were also controlled regarding the collocation frequencies between words and the frequencies of probing verbs with respect to their location (i.e., in main clauses or relative clauses), with half being correct/incorrect.

#### Verbal WM materials

The verbal WM task aimed to detect the linear memory ability without hierarchical processing, such that the stimuli in verbal WM task were generated matching the linear word sequential pattern of sentence processing stimuli (Chen et al., 2023; Liu et al., 2023; Zaccarella et al., 2017b). 6 nouns or 6 verbs were arranged in linear sequences to form a noun list or a verb list (Figure 2B). This task shared the same pool of words with the sentence processing task. A total of 36 noun lists and 36 verb lists were generated. The frequencies of word appearance and collocation were also controlled. The probing statement for word list trial was like “帮助*-5?*”, which asked participants to judge whether “帮助” appeared at the fifth position of the word list (Figure 2B). Half of the probing statements were correct/incorrect. The questioned words and their locations in the sequence were also balanced.

#### Visual WM materials

An orientation WM accuracy task was applied to assess the ability of visual working memory by requiring participants to remember the orientations of a set of items. The stimuli of the visual WM task were presented on the screen with a black fixation point surrounded by different number of bars (2° in length and 0.5° in width). All bars were presented within two 4° × 7.3° rectangular regions that were centered 3° to the left and right of the central fixation point (0.4° × 0.4°). Visual WM task consisted of two experimental conditions (load 2 and load 6) and a catch trial condition. For two experimental conditions, one or three bars were placed on each hemifield left or right to the fixation point for load 2 or load 6 condition. The orientation of bars was set at random between 0° and 180° but any two bars on the same screen were at least 20° apart (Figure 2C).

### 2.3 tPBM Protocol

The 1064-nm tPBM stimulation was conducted using a diode-pumped solid-state laser with a linewidth of ±1 nm (Model JL-LS-100 developed by Jieliang Medical Device Inc., Jiangxi, China). The 150 mW/cm^2^ power density dosage of the laser beam was adopted, with total area of 13.57 cm^2^, resulting a continuous power output of 2036 mW. The energy emitted by the laser diode at this setting was only 15% of the Maximum Permissible Exposure (MPE) to the skin (i.e., 1.0 W/cm^2^) according to the ANSI Z136.1-2014 standard, with no adverse effects detected from previous studies (Wang X. et al., 2022). The laser device was handheld, and participants were instructed to wear protective eyewear provided by the laser device manufacturer to protect their eyes from laser light. The stimulation site is shown by Figure 3A, the edge of which is along the eyebrow and hairline on the left forehead. In reference to the standard 10-20 EEG electrode placement system, the area stimulated roughly covered the ventral area of LPFC (i.e., LIFG). Both active and sham tPBM stimulation lasted for 16 min. No laser beam was emitted during sham sessions. The ambient noise (mainly caused by the cooling fan in the machine) was the same for sham or active sessions.

**Figure 3.**
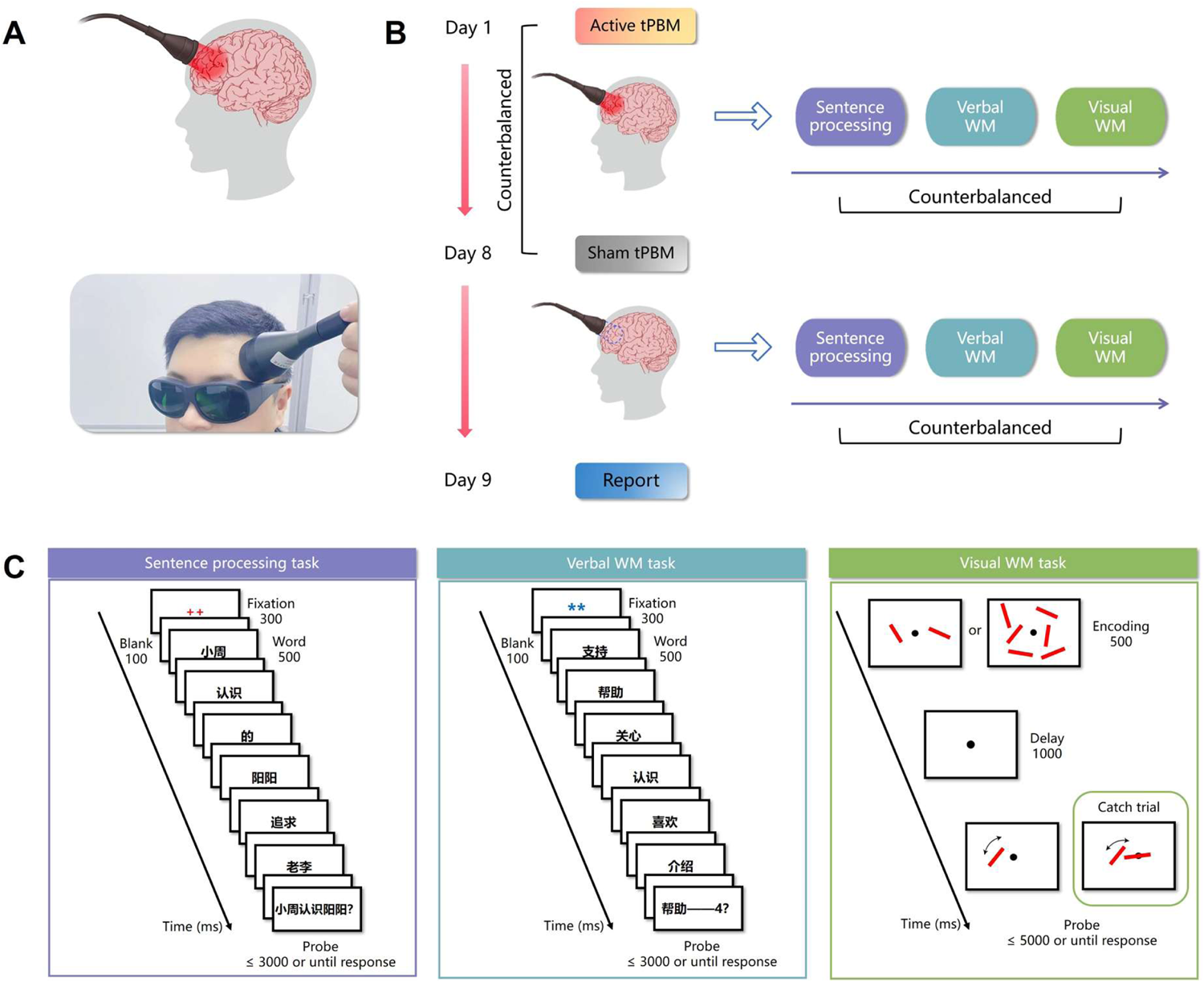
(**A**) The stimulation site was located to the ventroposterior part of left prefrontal cortex (LPFC) as shown in the diagram (upper) and the picture (lower) on which the 1064-nm tPBM was being applied to a simulated participant. (**B**) The experimental procedure of tPBM stimulation. Two sessions of tPBM were separated by seven days with one active and one sham tPBM session. After 16-min tPBM stimulation, three tasks were accomplished in counterbalanced order. At the 9^th^ day, participants were asked to report or guess in which session they received active tPBM stimulation according to their subjective feeling. **(C)** The procedures of three tasks. Sentence processing task and verbal WM task presented the trials word by word and asked participants to make T/F judgements on the probe screens. In the visual WM task, participants were asked to adjust the rotatable bar to its original position after encoding and delay screens. The catch trial presented a fixed bar across the fixation point and asked participants to turn the rotatable bar parallel to it.

### 2.4 Procedures

#### 2.4.1 Experimental procedure

The current study adopted a double-blind, sham-controlled tPBM experimental protocol. Specifically, each participant completed two experimental sessions separated by at least seven days to minimize the practice effect. One sham (placebo) stimulation session and one active stimulation session were performed respectively, the order of which was randomized and counterbalanced across participants (Figure 3B). The purpose and design of the current experiment were covered up towards participants.

At the beginning of each experimental session, participants performed training tasks first to ensure all of them could reach above the chance level of accuracy of all tasks. 16-min tPBM treatment was conducted then, during which participants were required to keep awake and mute. Three tasks were performed with counterbalanced order across participants immediately after the tPBM treatment. All participants reported no feelings or only minor feelings of tPBM treatment. At the day after the second session, participants were required to report or guess which session they thought to be the active stimulation session (Figure 3B). Results showed that participants guessed below the chance level (hit = 35.71%; miss = 32.14%; uncertain = 32.14%), suggesting that they were not aware of the condition of active or sham tPBM stimulation.

#### 2.4.2 Procedures of tasks

As for sentence processing task and verbal WM task, stimuli were presented through E-Prime version 3.0 (Psychology Software Tools, Inc., Pittsburgh, PA, USA). Specifically, complex sentences and word lists were presented word by word, with one word for 500 ms followed by a 100-ms blank. Attached to each sentence or word list, a probing statement were presented in whole sentences and lasted for a maximum of 3 s. Participants were instructed to judge the statement’s correctness and to press the corresponding buttons on the keyboard. The screen for probing statements terminated immediately after the participants press the button and was followed by a 1000-ms intertrial interval (ITI) (Figure 3C). Complex sentences were presented in a pseudorandom order [i.e., sentences of the same RC type (ORC or SRC) would not appear in more than three times consecutively]. Similarly, no more than three noun or verb word lists would appear consecutively in a pseudorandom order in verbal WM task.

In the visual WM task, the screen of memory encoding was presented for 500 ms and followed by a 1000-ms blank screen of delay. Next, the probing screen was presented for at most 5 seconds with a rotatable bar appearing at any position of two or six bars among the encoding arrays. Participants were instructed to adjust the bar with the mouse to the orientation according to their memory of the coded bars and press the left button. For the catch trial condition, a fixed bar with random orientation would lie across on the fixation point with another rotatable bar presented at a random place aside from the fixation point. Participants needed to adjust the orientation of the rotatable bar parallelly to the fixed bar in at most 5 seconds and press the left button of the mouse (Figure 3C). All screens of visual WM task were presented using Psychtoolbox version 3.0.19 (Kleiner et al., 2007) in Matlab version R2020b (MathWorks Inc., Natick, MA). The whole run of this task consisted of 5 blocks with 240 trials in total (96 of load 2, 96 of load 6, and 48 catch trials in random order). Participants could have a rest between two blocks.

### 2.5 Data Analyses

The data of accuracy (ACC) and response time (RT) of true/false judgement were directly recorded in sentence processing task and verbal WM task. As for the visual WM task, the differences between the real orientation of the bar and response orientation and RTs were collected firstly. To unify the dependent variable calculated from every task, the accuracy data was acquired further. Owing to the setting of 20° apart between any two bars, the response to one trial was classified as correct (i.e., ACC = 1) if the difference was between ±20°.

To avoid accuracy (ACC) - response time (RT) trade-offs, a measure of *overall performance* was used, which weighted the RT with the error rate (ER) according to the formula: *P* = *RT*(1 + 2*ER*), in which *ER* was equal to “1 – *ACC*” (Lyone et al., 2014). This measure could be interpreted as an adjusted RT penalized for inaccurate performances, where a higher value indicates worse performance (Lyons et al., 2014). The behavioral changes between sham and active conditions (i.e., the behavioral advantages brought by tPBM effect) were acquired by the differences of *P* (Δ*P = sham* – *active*) between the two stimulation conditions. The data of each group in the current research was interpolated according to the box plot. Outliers which were beyond *Q1* – 1.5\**IQR* or *Q3* + 1.5\**IQR* were interpolated by the values of *Q1* – 1.5\**IQR* or *Q3* + 1.5\**IQR* correspondingly. This method of data cleaning could reduce the effect of extreme values while keeping the data distribution relatively stable.

To certify the global effectiveness of tPBM modulation on the two groups, tests of 2-way mixed-effect repeated-measure analysis of variance (ANOVA) with stimulation condition (sham and active) and group (L1 and L2) as factors was performed on *P* for each task. Given the common practice to group participants depending on high and low cognitive capacities in neuromodulation studies, which usually found that cognitive ability improvement existed mainly or more robustly for individuals with lower original capacity (Hsu et al., 2014; Tseng et al., 2012), the analyses with the same purpose were conducted in the current study in case tPBM showed a significant enhancement. Nevertheless, we did not simply group participants in subgroups of low and high primal capacity considering the fact that the proportion of orders of tPBM sessions the participants were assigned with (i.e., sham stimulation first or active stimulation first) could be unbalanced in different subgroups. A correlation analysis between initial performance (i.e., *P* on sham condition) and the change of performance (Δ*P*) was performed instead, which could certify the correlation if the lower initial performance correlated larger change of performance after tPBM stimulation.

The statistical tests in the current study were accomplished through JASP version 0.17.2.1 (https://jasp-stats.org/) and R version 4.3.1 (R Foundation for Statistical Computing, Vienna, Austria; https://www.R-project.org/).

## 3 Results

### 3.1 tPBM over LPFC enhanced sentence processing performance in both L1 and L2 participants

For sentence processing task, the results of 2-way mixed-effect ANOVA (Figure 4A) showed a significant main effect of stimulation condition [*F* (1, 54) = 10.931, *p* = 0.002, η_p_^2^ = 0.168]. Specifically, compared with sham tPBM stimulation condition, the performance on the active session was significantly better with lower *P* value (sham: 2556.09 ± 733.21 ms; active: 2343.36 ± 674.59 ms), suggesting the increased performances of complex sentence processing both for L1 and L2 group due to tPBM. The follow-up paired sample *t* test revealed that the active tPBM enhanced sentence processing performance for both L1 [*t* (27) = 2.085, *p* = 0.047, Cohen’s *d* = 0.394; sham = 2104.35 ± 442.42 ms, active: 1948.92 ± 395.06 ms] and L2 [*t*(27) = 2.575, *p* = 0.016, Cohen’s *d* = 0.487; sham = 3009.46 ± 688.90 ms, active: 2737.80 ± 669.50 ms] groups. In addition, the ANOVA also manifested the strong main effect of group factor [*F* (1, 54) = 38.592, *p* < 0.001, η_p_^2^ = 0.417], such that L1’s performance (2026.63 ± 422.91 ms) was much better than L2 (2873.63 ± 686.88 ms), suggesting the different ability with regard to language proficiency. Moreover, the null result of interaction effect of ANOVA [*F* (1, 54) = 0.809, *p* = 0.372, η_p_^2^ = 0.015] revealed that L1 and L2 groups showed similar extent to be enhanced by tPBM.

**Figure 4.**
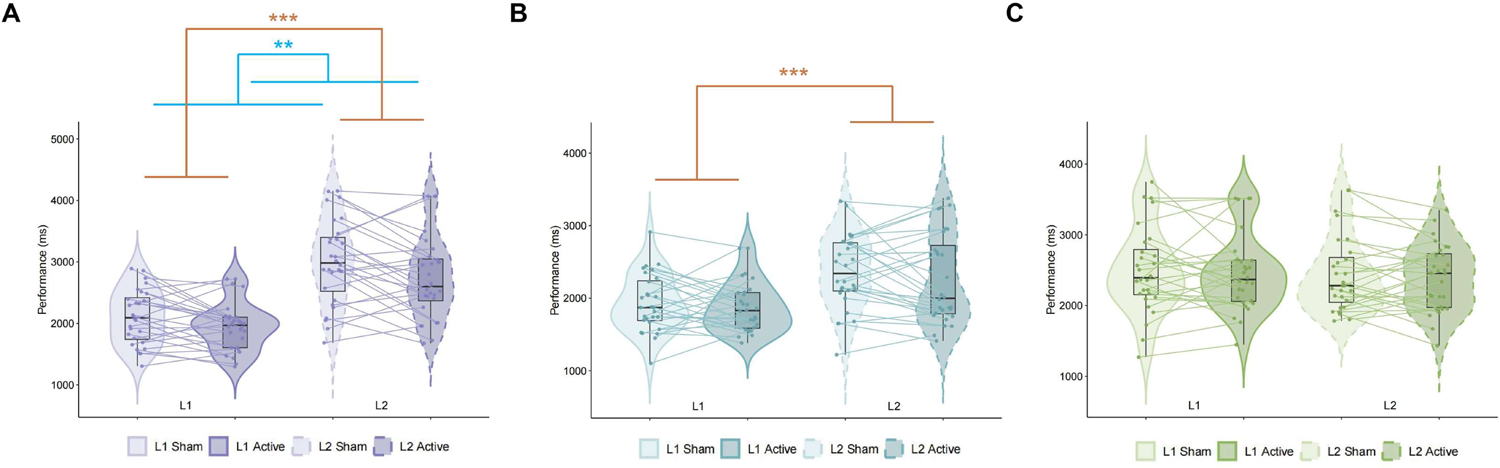
The violin plots of ANOVA results in (**A**) sentence processing task; (**B**) verbal WM task; (**C**) visual WM task. Each dot refers to one participant. The lines connect the measurements of the same individuals. The line in the box plot represents the median of the data. The violin plots for L1 and L2 are bordered with solid and dashed lines respectively. The plots in darker color refers to sham condition and the lighter one refers to the active condition. The blue line of significance shows the main effect of stimulation condition and the orange one shows the main effect of group. * *p* < 0.05; ** *p* < 0.01; *** *p* < 0.001.

### 3.2 tPBM over LPFC failed to enhance working memory performance

Similarly, a 2-way mixed-effect ANOVA was performed with stimulation condition (sham and active) and group (L1 and L2) as factors on the verbal working memory task (Figure 4B). However, no stimulation condition main effect [*F* (1, 54) = 1.835, *p* = 0.181, η_p_^2^ = 0.033] or interaction effect [*F* (1,54) = 0.223, *p* = 0.639, η_p_^2^ = 0.004] between group and stimulation condition could be identified. This result suggested that active tPBM on LPFC (esp., LIFG) did not benefit the performance on verbal working memory as opposed to sentence processing for both L1 and L2 groups. Considering the discrepant coding difficulty toward Chinese words between L1 and L2, the main effect of group was pronounced [*F* (1, 54) = 12.914, *p* < 0.001, η_p_^2^ = 0.193], such that L1 reached better performance (1903.58 ± 371.30 ms) with lower *P* value when compared to L2 (2312.63 ± 564.48 ms).

As for the visual working memory task which was manipulated as a non-language control in the current study, a 2-way mixed-effect ANOVA showed null effects either for group main effect [*F*(1, 54) = 0.549, *p* = 0.462, η_p_^2^ = 0.004], stimulation condition main effect [*F* (1, 54) = 0.016, *p* = 0.899, η_p_^2^ = 0.010], or the interaction between them [*F* (1, 54) = 0.234, *p* = 0.631, η_p_^2^ < 0.001] (Figure 4C) as expected. The current results revealed the fact that the tPBM stimulation on LPFC exerted little effect on visual working memory regardless of groups of participants. Furthermore, the two groups showed similar performance on visual working memory task in contrast to the two language-related tasks above.

### 3.3 Inability to process complex sentences predicts large tPBM benefits

To test whether the extent of tPBM boost related to the primal performance in sentence processing task, a correlation task between *P* on sham condition and Δ*P* was conducted. Given that the initial performances between sentence processing task and the other two tasks were highly correlated (sentence processing & verbal WM tasks: Pearson’s correlation *r* = 0.700, *p* < 0.001, Figure 5A; sentence processing & visual WM tasks: Pearson’s correlation *r* = 0.281, *p* = 0.036, Figure 5B), the Pearson correlation analysis between initial performance (i.e., *P* on sham condition) and the change of performance (Δ*P*) was conducted with initial performance of verbal working memory task and visual working memory task partially out, which could provide the reliable evidence of the correlation. As expected, the initial performance on the sentence processing task was positively correlated with the chance of performance (sham - active) on the same task (Pearson’s correlation *r_partial_* = 0.384, *p* = 0.004; Figure 5C).

**Figure 5.**
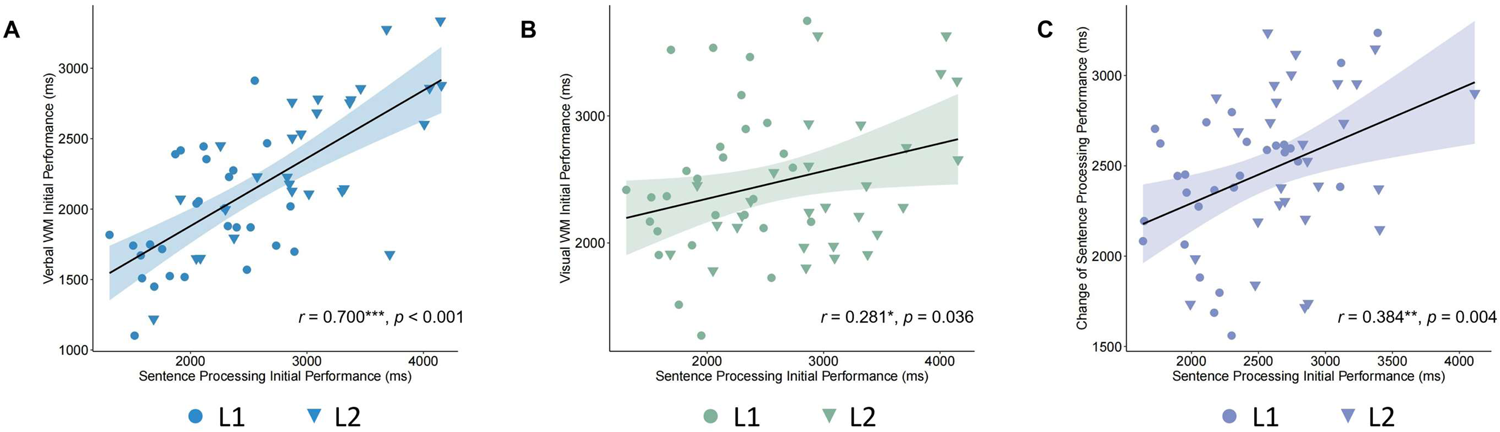
The correlation between (**A**) Sentence processing initial performance (i.e., *P* on sham condition) & verbal working memory (WM) initial performance; (**B**) Sentence processing initial performance & visual WM initial performance; (**C**) Sentence processing initial performance & change of sentence processing performance. The shaded areas represent 95% confidence intervals. Circles and triangles represent L1 and L2 participants respectively. * *p* < 0.05; ** *p* < 0.01; *** *p* < 0.001.

## 4 Discussion

In the present study, we mainly applied tPBM to L1 and L2 sentence processing task(s) with verbal WM task and visual WM task additionally involved, aiming to investigate the tPBM effect on sentence processing and figure out the possible effective pattern interfered by WM in L1 speakers and L2 learners. Results showed that tPBM on LPFC selectively enhanced the sentence processing performances rather than the WM task performances reflecting the verbal or the visual WM capacities in both L1 and L2 groups, and the current results did not support an interfering role of verbal WM in-between the tPBM stimulation and the modulation on sentence processing. In sentence processing task, making judgments on probing statements of thematic role assignment required reordering and integration of sentential elements (Friederici, 2017; Xu et al., 2020), thereby the overall performance (indicator combining the ACC and RT) of sentence processing task could reliably reflect the sentence processing ability. Together, our results supported the direct effective pathway of tPBM effect on sentence processing performances both for L1 and L2 speakers and demonstrated that tPBM could enhance sentence processing ability without the interference of WM. The null results of the interaction effect between the group and stimulation type factor validated that L1 and L2 showed similar patterns of modulation. Moreover, the non-significant results of tPBM on WM capacities suggested that tPBM on LPFC (esp., the ventral part of LPFC covering the LIFG) was specific to language (esp., complex sentence) processing. Specifically, L1 and L2 differed in language-related tasks (sentence processing task and verbal WM task) but not in the nonverbal task of visual WM, indicating that L1 and L2 matched on nonverbal task so that the parallel pattern of modulation between L1 and L2 was consolidated. In the sentence processing task, we further found that participants with worse initial performance received more enhancement through tPBM such that the inability to process complex sentence can predict large tPBM benefits, which is consistent with the results from prior neuromodulation studies (Hsu et al., 2014; Tseng et al., 2012).

### 4.1 The effect of tPBM on sentence processing ability without WM’s interference

With converging evidences showing that tPBM reveals enhancement towards cognitive abilities such as WM, attention, and executive functions, tPBM has been acknowledged as a promising NIBS technique for neuromodulation (Barrett & Gonzalez-lima, 2013; Blanco et al., 2017; Chan et al., 2019, 2021; Holmes et al., 2019; Hwang et al., 2016; Moghadam et al., 2017; Qu Xiu. et al., 2022; Zhao et al., 2022). The current study not only extended tPBM’s application to the high-level cognitive ability specific to human beings—sentence processing ability by using the sentence processing task in L1 and L2 groups but also scrutinized the relationships between language and WM capacities on the basis of neuromodulation. In the sentence processing task, participants needed to reorder sentential elements in RCs with syntactic movement and then to construct hierarchical structures, which cost a high load of syntactic computation (Friederici, 2017; Xu et al., 2020). Combined with our results indicating that the ability of sentence processing through syntactic merge operation could be significantly enhanced, it became a novel complementary finding that in general, the metabolic and hemodynamic changes induced by tPBM on LPFC could also boost one of the highest-level cognitions of human-beings—language faculty. Furthermore, our results showed that tPBM only came into effect in sentence processing task instead of WM tasks, which revealed the direct pathway of tPBM effect on sentence processing ability with no potential interference of WM. To note, the selective tPBM enhancement pattern did not deny the contribution of verbal WM ability to sentence processing as discussed to a large extent in the prior relative studies (Caplan & Waters, 1999; Grossman et al., 2002; Makuuchi et al., 2009; Meyer et al., 2012; Santi & Grodzinsky, 2007). Nevertheless, the present study revealed that tPBM could enhance sentence processing ability directly through stimulating LPFC, thus providing reliable evidence that such an enhancement should not be caused by a general increase of the verbal working memory capacity as a by-product. Hence, this study is in support of the functionally-specific role of the ventral LPFC (esp., LIFG) on language/sentence processing.

It has already been proposed that LPFC is actively involved in different forms of hierarchical processing, in which BA 44 in LIFG may play an integral and essential role in the process (Jeon, 2014). LIFG was further delineated as the region responsible for the merge operation of sentence processing in several recent neurolinguistic studies, in which sentence processing was compared to word list (where syntax is subtracted away) processing in order to purify the neural basis of merge (Snijders et al., 2009; Zaccarella et al., 2017a; Wu et al., 2019). Meanwhile, the LIFG’s engagement was found not only in inflecting languages like German (Zaccarella et al., 2017a) or Dutch (Snijders et al., 2009), but also in Chinese and other languages devoid of morphological inflections (Chen et al., 2023; Wu et al., 2019). For instance, Wu et al. (2019) compared the two-word Chinese phrase consisting of a determiner and a classifier to the two-word list consisting of two classifiers and found that LIFG (specifically BA 44 and BA 45) was significantly activated in the process of phrase building. These findings converged on the notion that LIFG’s engagement in sentence processing was cross-lingual (Chen et al., 2023; Wu et al., 2019). Along with natural language, artificial grammar was also exploited to investigate the neural basis of the hierarchical building, through which the semantic confounders could be excluded and all critical variables could be better controlled across the participants (Friederici, 2011; Gómez & Gerken, 2000; Jeon, 2014; Uddén & Männel, 2018). Similarly, studies with diverse types of artificial grammars also pinpointed that LIFG is working as a combinatorial engine where words were merged together and sentences were built (e.g., Bahlmann et al., 2008; Chen et al., 2021a; Friederici et al., 2006a; Liu et al., 2023). Furthermore, studies of NIBS provided us with causal evidence of the relationship between LIFG and sentence processing ability (de Vries et al., 2010; Kuhnke et al., 2017; Meyer et al., 2018; Sakai et al., 2002; Uddén et al., 2017). TMS was adopted in Kuhnke et al. (2017) to suggest the causal involvement of LIFG in reordering during sentence processing. De Vries et al. (2010) found Broca’s area (BA44/45 in LPFC) was causally related to the ability to detect syntactic violations in artificial grammar by means of transcranial direct current stimulation (tDCS). Given that the stimulation site of tPBM of the present study is on the ventral part of LPFC, mainly covering the LIFG, our results provided new evidence for a causal role of LPFC (esp., LIFG) for sentence processing (possibly complex syntactic processes), through the positive intervention effect of tPBM for the first time.

### 4.2 L1 and L2 participants showed similar pattern of modulation

One of our most interested research questions is whether L2 learners could exhibit a similar tPBM enhancement pattern on sentence processing as in L1 speakers. The current study found that L2 showed a similar pattern of modulation with L1 speakers, with sentence processing ability improved after tPBM stimulation but not for WM tasks. These results are in line with the notion that sentence processing in L1 and L2 both involved LIFG. Studies have shown that LIFG plays an integral part in L2 sentence processing and yields large plasticity (e.g., Chen et al., 2019, 2021b, 2023; de Vries et al., 2010; Luke et al., 2002; Nakagawa et al., 2022; Wartenburger et al., 2003). Results from studies adopting natural language materials converged on the fact that LIFG was required in the course of sentence processing and learning (Luke et al., 2002; Musso et al., 2003; Nakagawa et al., 2022; Sakai et al., 2004; Wartenburger et al., 2003; Yusa et al., 2011). Specifically, Nakagawa et al. (2022) pinpointed the role of LIFG for hierarchical syntactic processing by involving English L2 learners whose native language was Japanese. A study of L2 learning from Sakai et al. (2004) substantiated the neural plasticity by showing that the activation of LIFG was boosted after 2-month L2 (English) training and practicing. The evidence from late L2 learners even found activation of LPFC to a higher extent than their L1 processing when participants read L2 sentences, showing that lower language proficiency led to higher brain calling (Luke et al. 2002; Wartenburger et al., 2003). The artificial grammar learning paradigm provided us with more insights into the neural basis of L2 syntactic learning (Bahlmann et al., 2008; Chen et al., 2019, 2021a, 2021b, 2023; Friederici et al., 2006a; Grey et al., 2018; Liu et al., 2023; Morgan-Short et al., 2015). Chen et al. (2019, 2021b) proposed that L2’s sentence structure learning based on word category information involved posterior LIFG. A novel artificial hierarchical syntactic structure-building grammar was developed by Chen et al. (2021a) and Liu et al. (2023), and demonstrated that the fundamental operation of merge rested on the function of BA 44 in LIFG. To sum up, sentence processing activates LIFG both for L1 and L2 reading, which shows large plasticity to be modulated, although some studies pointed out that L2 processing might involve more anterior regions in LIFG with lower automaticity (Jeon & Friederici, 2013).

### 4.3 Application prospect of tPBM on sentence processing

Consistent with former studies of neuromodulation (Hsu et al., 2014; Tseng et al., 2012), the current study found that participants with lower sentence processing ability at the initial state (i.e., prior to tPBM) were more susceptible of tPBM improvement. State-dependency was often used to explain the effects of brain stimulation and the initial state of stimulated regions (Hsu et al., 2014; Silvanto et al., 2008). For instance, TMS has been shown to be particularly effective on neurons that are less active (Silvanto et al., 2008). As for the current study, the state-dependent effect of neuromodulation techniques may become a feasible interpretation also for tPBM, that is, lower performers may have neurons less activated initially and thus show greater tPBM effects in return.

More importantly, the correlation between initial sentence processing performance and the degree of tPBM improvement broadened the prospect of tPBM applications. With the fact that tPBM was more effective for lower performers, the value of tPBM became more prominent by applying tPBM towards less-competent groups. Furthermore, we certified the positive modulation effect of tPBM on L2 learners, who served as the participants with lower (L2) complex sentence processing ability. The present findings, therefore, suggested that it might be available to further apply tPBM to the upregulation of participants with language ability deficiency. Overall, the present study shed light on tPBM—a promising NIBS tool/approach— for its future clinical applications on the population struggling with language acquisition/learning difficulties, language impairments, or progressing language capacity declination. In the future, tPBM is expected to be a favorable alternative with relatively low cost and highly consistent enhancement effect to improve/facilitate the human language faculty.

### 4.4 Limitations

Although LIFG was acknowledged to play a key role in hierarchical syntactic structure construction, sentence processing also involves several other crucial brain regions such as left posterior temporal cortex (LpTC; Chen et al., 2023; Kinno et al., 2008) and is supported by a left-dominant fronto-temporal network (Friederici, 2017). The current study only investigated the tPBM effect through LPFC and did not go deeper into the intervention on brain network, which was worth exploring further. Moreover, syntactically-complex sentence processing is also accompanied with the difficulty of semantic interpretation, and given the comparatively coarse stimulation location of tPBM, the enhancement effect of sentence processing might be related to the facilitation of both syntactic and semantic processes. Although the present study treated the sentence processing as a holistic/unified process, future studies might employ jabberwocky sentences (e.g., Friederici et al., 2006a; Matchin et al., 2019; Zhang et al., 2022) or artificial grammars (e.g., Bahlmann et al., 2008; Chen et al., 2021a; Liu et al., 2023) to further differentiate these internal linguistic processes.

Moreover, the neural mechanisms underlying the tPBM effects on behavioral performances of language/sentence processing still remains unclear in the present. Further studies are expected to provide neuroimaging data and make further exploration and interpretation of the neural changes brought by tPBM.

## 5 Conclusion

The present study applied the novel NIBS technique—tPBM on LPFC to upregulate the sentence processing performances. As shown by the behavioral performance changes, tPBM improved the sentence processing ability in both L1 and L2 groups. Moreover, L1 and L2 participants showed consistent tPBM enhancement pattern without the interference of verbal WM. It is also noteworthy that participants with lower initial sentence processing performances would benefit more from tPBM. Taking together, these findings supported the positive effectiveness of tPBM on high-level human cognitions and unprecedentedly extended tPBM’s application to human language faculty as reflected by complex sentence processing performances, and thus, such a promising and cost-effective NIBS tool is of great social and clinical significances for future applications.

## CRediT authorship contribution statement

**Mingchuan Yang & Yang Liu:** Methodology, Data curation, Formal analyses, Visualization, Writing – original draft. **Zhaoqian Yue:** Data curation, Writing – review & editing. **Guang Yang, Xu Jiang, Yimin Cai, & Yuqi Zhang:** Writing – review & editing. **Xiujie Yang**: Supervision, Writing – review & editing. **Dongwei Li:** Conceptualization, Methodology, Supervision, Writing – review & editing. **Luyao Chen:** Conceptualization, Methodology, Supervision, Writing – review & editing, Funding acquisition.

## Data availability

The data, code, and materials of this study are available at https://osf.io/e35ac/. Further data and materials which also form other ongoing studies will be made available upon reasonable requests and collaborative agreement addressed to the co-authors by contacting the Max Planck Partner Group, School of International Chinese Language Education, Beijing Normal University, and signing a formal data-sharing agreement.

## Acknowledgments

This work was supported by the National Social Science Foundation of China (No. 22CYY017). The authors wish to thank all participants who took part in this study. Special thanks are extended to Xuye Yuan and Yitong Wu for the assistance on data collection.

## Footnotes

1 The control of L2 learners’ mother tongues aimed to increase the typological differences between Chinese and their L1s so that they could process L2 sentences in a distinctive fashion from L1, which could minish the confounding effect brought by the syntactic similarity.

